# Devil in the details: growth, productivity, and extinction risk of a data-sparse devil ray

**DOI:** 10.1101/043885

**Authors:** Sebastián A. Pardo, Holly K. Kindsvater, Elizabeth Cuevas-Zimbrón, Oscar Sosa-Nishizaki, Juan Carlos Pérez-Jiménez, Nicholas K. Dulvy

## Abstract

Devil rays (*Mobula* spp.) face rapidly intensifying fishing pressure to meet the ongoing international trade and demand for their gill plates. This has been exacerbated by trade regulation of manta ray gill plates following their 2014 CITES listing. Furthermore, the paucity of information on growth, mortality, and fishing effort for devil rays make quantifying population growth rates and extinction risk challenging. Here, we use a published size-at-age dataset for a large-bodied devil ray species, the Spinetail Devil Ray (*Mobula japanica*), to estimate somatic growth rates, age at maturity, maximum age and natural and fishing mortality. From these estimates, we go on to calculate a plausible distribution of the maximum intrinsic population growth rate (*r*_*max*_) and place the productivity of this large devil ray in context by comparing it to 95 other chondrichthyan species. We find evidence that larger devil rays have low somatic growth rate, low annual reproductive output, and low maximum population growth rates, suggesting they have low productivity. Devil ray maximum intrinsic population growth rate (*r*_*max*_) is very similar to that of manta rays, indicating devil rays can potentially be driven to local extinction at low levels of fishing mortality. We show that fishing rates of a small-scale artisanal Mexican fishery were up to three times greater than the natural mortality rate, and twice as high as our estimate of *r*_*max*_, and therefore unsustainable. Our approach can be applied to assess the limits of fishing and extinction risk of any species with indeterminate growth, even with sparse size-at-age data.

## Introduction

Understanding the sustainability and extinction risk of data-sparse species is a pressing problem for policy-makers and managers. This challenge can be compounded by economic, social and environmental actions, as in the case of the mobulid rays (subfamily Mobulinae). This group includes two species of charismatic and relatively well-studied manta rays (*Manta* spp.), which support a circumtropical dive tourism industry with an estimated worth of #73 million USD per year [1]. The Mobulinae also includes nine described species of devil rays (*Mobula* spp.). The recent international trade regulation of manta ray gill plates under the Convention of International Trade of Endangered Species (CITES) [2] is likely to shift gill plate demand from manta rays onto devil rays.

Devil rays are increasingly threatened by target and incidental capture in a wide range of fisheries, from small-scale artisanal to industrial trawl and purse seine fisheries targeting pelagic fishes [3,4]. The meat is sold on domestic markets and the gill plates are exported to meet consumer demand, mostly from China [5]. Small-scale subsistence and artisanal fisheries, mainly for meat, have operated throughout the world for decades [5]. For example, devil rays were caught by artisanal fishermen using harpoons and gill nets around Bahia de La Ventana, Baja California, Mexico, until 2007 when the Mexican government prohibited the take of mobulid rays [6].

Overall, around 90,000 devil rays are estimated to be caught annually in fisheries worldwide [7]. Many industrial fleets capture devil rays incidentally. For example, European pelagic trawlers in the Atlantic catch a range of megafauna including large devil rays at a rate of up to one individual per hour [3], while purse seine fleets targeting tunas capture tens of thousands of devil rays each year [4]. Even if devil rays are handled carefully and released, their post-release mortality might be significant [8]. We do not know whether the fisheries and international trade demand for devil rays are significant enough to cause population declines and potential extinction. The degree to which devil ray populations can withstand current patterns and levels of fishing mortality depends on their intrinsic productivity, which determines their capacity to compensate for fishing.

Slow somatic growth and large body size are associated with low productivity and elevated threat status and extinction risk in marine fishes, including elasmobranchs [9–11]. Based on these correlations, the American Fisheries Society developed criteria to define productivity and extinction risk: They defined four levels of productivity (very low, low, medium, and high) based on four life history traits (age at maturity, longevity, fecundity, and growth rate, which is related to the von Bertalanffy growth coefficient *k*) and the intrinsic rate of population increase *r* [9]. According to these criteria, manta rays have very low or low productivity, with some of the lowest maximum rates of population increase (*r*_*max*_) of any shallow-water chondrichthyan [9,12].

Here we evaluate the productivity, and hence relative extinction risk of large devils rays, using the only age and growth study available for this group [13]. We use a Bayesian estimate of somatic growth rate and a demographic model based on the Euler-Lotka equation to calculate the maximum intrinsic rate of population increase (*r*_*max*_) for a population of the Spinetail Devil Ray *Mobula japanica* (Müller & Henle, 1841) and compare it to the productivity of 95 other sharks, rays, and chimaeras.

## Methods

We take advantage of the first study to measure length-at-age for catch data of *M. japanica* [13]. The Spinetail Devil Ray is similar in life history and size to other exploited mobulids, so we assume it is representative of the relative risk of the group. Spinetail Devil Rays examined in this study were caught seasonally by artisanal fishers using harpoon and gill nets around Punta Arenas de la Ventana, Baja California Sur, Mexico, during the summers of 2002, 2004, and 2005 [13].

First, we estimate growth parameters using a Bayesian approach that incorporates prior knowledge of maximum size and size at birth of this species, using the length-at-age data presented in Cuevas-Zimbrón et al. (2013) [13] (Part 1). Second, we use the same dataset to plot a catch curve of the relative frequency of individuals in each age-class, from which we can infer a total mortality rate (*Z*) that includes both fishing (*F*) and natural mortality (*M*) (Part 2). This places an upper bound on our estimate of natural mortality, and allows us to compare the observed rate of mortality for this population with independent estimates of natural mortality rates. Third, we estimate the maximum intrinsic rate of population increase (*r*_max_) for this devil ray (Part 3) and compare it against the *r*_*max*_ of 95 other chondrichthyans, calculated using the same method (Part 4).

### Part 1: Re-estimating von Bertalanffy growth parameters for the Spinetail Devil Ray

We analyse a unique set of length-at-age data for a single population of *M. japanica* caught in a Mexican artisanal fishery. Individuals in this sample were limited to 110 and 240 cm disc width (DW), which falls short (77%) of the maximum disc width reported elsewhere [14]. Therefore, we use a Bayesian approach to refit growth curves to this length-at-age dataset [15]. We use published estimates of maximum size and size at birth to set informative priors.

We fit the three-parameter von Bertalanffy equation to the length-at-age data, combining sexes:

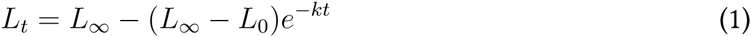

where *L*_*t*_ is length at age *t*, and the growth coefficient *k*, size-at-age zero *L*_0_, and asymptotic size *L*_∞_ are the von Bertalanffy growth parameters. These parameters are conventionally presented in terms of length; here we use length synonymously with disc width, such that *L*_*t*_, *L*_0_, *L*_∞_, and *L*_*max*_ are synonymous with *DW*_*t*_, *DW*_0_, *DW*_∞_, and *DW*_*max*_, respectively.

In order to account for multiplicative error, we log-transformed the von Bertalanffy growth equation and added an error term:

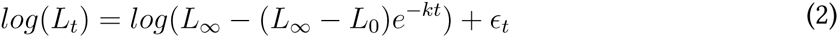

This can be written as:

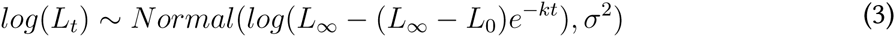

A Bayesian approach allows us to incorporate expert knowledge using prior distributions of estimated parameters. We based our informative priors on our knowledge of maximum disc widths and size-at-birth of *M. japanica*. Reported size at birth ranges from 88 to 93 cm DW [16,17], while reported maximum size for *M. japanica* is 310 cm DW [18]. While this reported maximum size is from an individual recorded in New Zealand, we use this estimate as genetic evidence suggests that populations of Spinetail Devil Rays in the Pacific Ocean have little genetic substructure [19]. Asymptotic size can be estimated from maximum size in fishes using the following equation [20]:

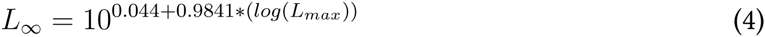

where *L*_*max*_ is maximum size, in centimetres. This results in an estimate of *L*_∞_ = 1.01 * *L*_*max*_ for a value of *L*_*max*_ = 310 cm DW. Instead of setting a fixed value for the conversion parameter, we create a hyperprior for this parameter, defined as *kappa*, based on a gamma distribution around a mean of 1.01. We concentrated the probability distribution of *kappa* between 0.9 and 1.1, and fully constrain it between 0.7 and 1.3 [20]:

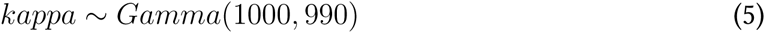

We also constrain our prior for *L*_*0*_ around size at birth, and use a beta distribution to constrain our prior for growth coefficient *k* between zero and one, with a probability distribution that is slightly higher closer to a value of 0.1:

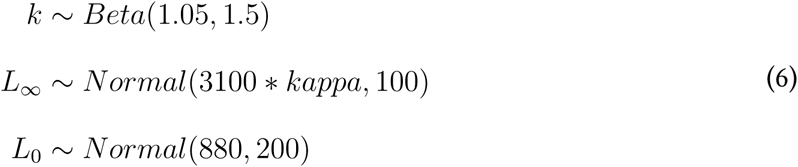

We compared the effect of our informative priors on our posteriors with parameter estimates with weaker priors, in which we maintained the mean of the distributions but increased their variance:

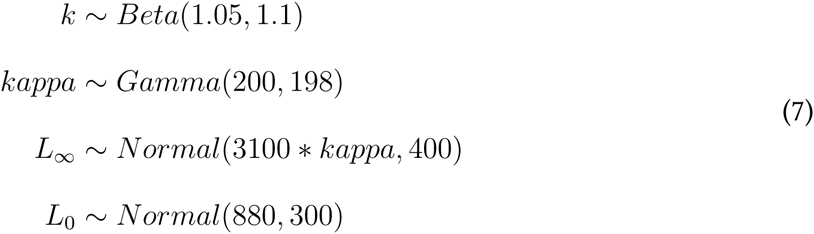

We also considered a scenario with uninformative priors, where all prior distributions are uniform:

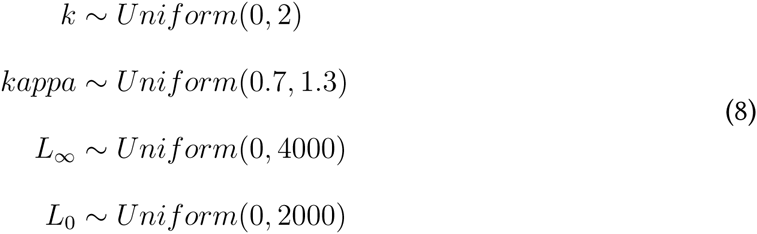

In all models we set an weakly informative prior for the variance *σ*^2^, such that:

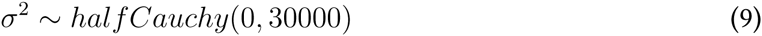

A summary of the priors used can be seen in Table 1. Bayesian inference was conducted using RStan v2.7.0 [21,22] running in R v3.2.1 [23].

**Table 1:**
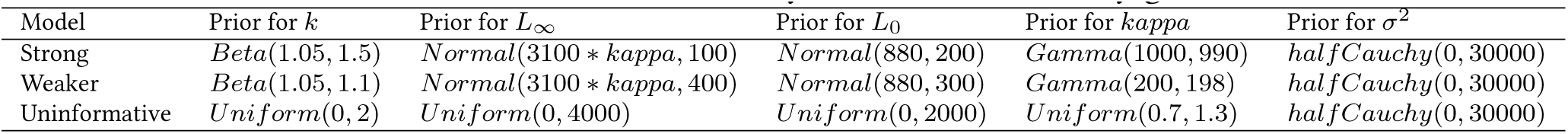
Priors used in the three di.erent Bayesian von Bertalan.y growth models.

### Part 2. Estimating total mortality using the catch curve

The length-at-age dataset of *M. japanica* can be used as a representative sample of the number of individuals within each age-class if we assume that sampling was opportunistic, and non-selective across each age‐ or size-class. We also assume that there is limited migration in and out of this population. With these assumptions, counting the number of individuals captured in each age-class represents the population age structure, which can be used to construct a catch curve.

Catch curves are especially useful for data-poor species lacking stock assessments [24,25]. The frequency of individuals in older or larger classes decreases due to a combination of natural and fishing mortality. If fishing is non-selective with respect to size, the total mortality rate *Z*, which is a combination of both fishing mortality *F* and natural mortality *M*, can be estimated using a linear regression as the slope of the natural log of the number of individuals in each class [26]. This information is very valuable when inferring whether fishing mortality *F* is unsustainable. We calculate Z as the slope of the regression of the catch curve, including only those ages or sizes that are vulnerable to the fishery.

We removed age-classes that had zero individuals in our sample to be able to take the natural logarithm of the count. Because there is uncertainty associated with the dataset (due to its relatively small size), we resampled a subset of the dataset 20,000 times, after randomly removing 20% of the points. This allowed us to quantify uncertainty in our estimate of *Z*. For each subset, we computed the age-class with the maximum number of samples, and removed all age-classes younger than this peak. With the remaining age-classes, we fit a linear regression to estimate the slope which is equivalent to – *Z*. This method for estimating mortality relies on two assumptions of the selectivity of the fishery. First, catch is not size-selective once individuals are vulnerable to the fishery. Second, if young age-classes are less abundant than older age-classes, they are assumed to have lower catchability. This is why we removed the younger age-classes before the “peak” abundance of each sample, as this will affect the steepness of the slope.

### Part 3. Estimating *M. japanica* maximum population growth rate

Maximum intrinsic population growth rates *r*_*max*_ can be estimated based on a simplified version of the Euler-Lotka equation [27, 28]. We use the following version of this equation to calculate *r*_*max*_. Unlike previous estimates of *r*_*max*_ for chondrichthyan species [12, 29, 30], this equation accounts for juvenile mortality [31]:

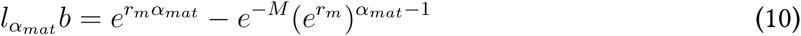

where *l*_*α*_*mat*__ is survival to maturity and is calculated as *l*_*α*_*mat*__ = (*e*^‒*M*^)^*α*_*mat*_^, *b* is the annual reproductive output of daughters, *α*_*mat*_ is age at maturity in years, and *M* is the instantaneous natural mortality. We then solve equation 10 for *r*_*max*_ using the `nlm.imb` function in R. To account for uncertainty in input parameters, we use a Monte Carlo approach and draw values from parameter distributions to obtain 10,000 estimates of *r*_*max*_. Next we describe how we determined each parameter distribution.

#### Annual reproductive output (*b*)

Adult female mobulid rays only have one active ovary and uterus where a single pup grows. This sets the upper bound of annual fecundity to one pup per year [14], and assuming a 1:1 sex ratio, results in an estimate of *b* of 0.5 female pups per year. It is possible female devil rays have a biennial cycle of reproduction where one pup is produced every two years, as in manta rays [12], so the lower bound for our estimate of *b* is 0.25 female pups per year. Thus we draw b from a uniform distribution bound between 0.25 and 0.5.

#### Age at maturity (*α*_*mat*_)

There are no direct estimates of age at maturity for any mobulid ray, but using age and growth data from Cuevas-Zimbrón (2013) [13] and a size at maturity of 200 cm DW from Serrano-López (2009) [32], we assume *M. japanica* individuals reach sexual maturity after 5-6 years. Thus we draw *α*_*mat*_ from a uniform distribution bound between 5 and 6.

#### Natural mortality (*M*)

We estimate natural mortality as the reciprocal of average lifespan: *M* = 1/*ω* where average lifespan *ω* is (*α*_*mat*_ + *α*_*max*_)/2). We used this estimate of *M* to calculate survival to maturity *l*_*α*_*mat*__ as (*e*^‒*M*^)^*α*_*mat*_^. This method produces realistic estimates of *r*_*max*_ when accounting for survival to maturity [31]. We also use our estimate of *Z* from Part 2, which represents both natural and fishing mortality, to contextualize our estimate of *M*. More specifically, our estimate of *Z* sets the upper bound for our estimate of *M*. We calculated maximum age (*α*_*max*_) based on the results of our analysis in Part 1. We therefore calculate *M* iteratively by drawing values of *α*_*mat*_ (described in the section above) and *α*_*max*_ from uniform distributions bound between the ranges mentioned.

### Part 4. Comparison of Mobula *r*_*max*_ among chondrichthyans

We re-estimate *r*_*max*_ for the 94 chondrichthyans with complete life history data examined in [12, 30, 31] using equation 10. We also update estimates of *r*_*max*_ for manta rays (*Manta* spp.) from Dulvy et al. 2014 [12], as a comparison with a closely related species.

## Results

### Part 1: Re-fitting the growth curve for *Mobula japanica*

The Bayesian model with strong priors yielded a lower estimate of *k* (0.12 year^−1^) and a higher estimate of *L*_∞_ (2995 cm DW) than the estimates based on weaker and uninformative priors (Table 2, Fig. 1). The asymptotic size in the model with strong priors was closest to the maximum observed size for this species (Fig. 2). Estimates of *k* were lowest in the model with strong priors and highest in the model with uninformative priors (Table 2, Fig. 1).

**Table 2:**
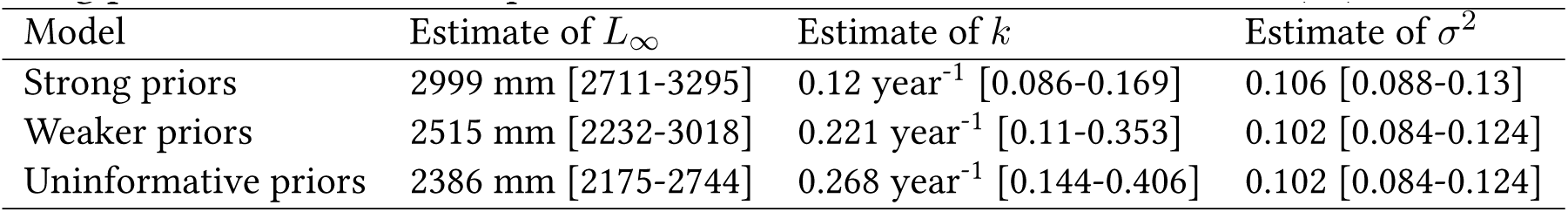
Mean von Bertalanffy growth parameter estimates for the three Bayesian models with differing priors. Values inside square brackets are the 95% credible intervals (CI).

**Figure 1:**
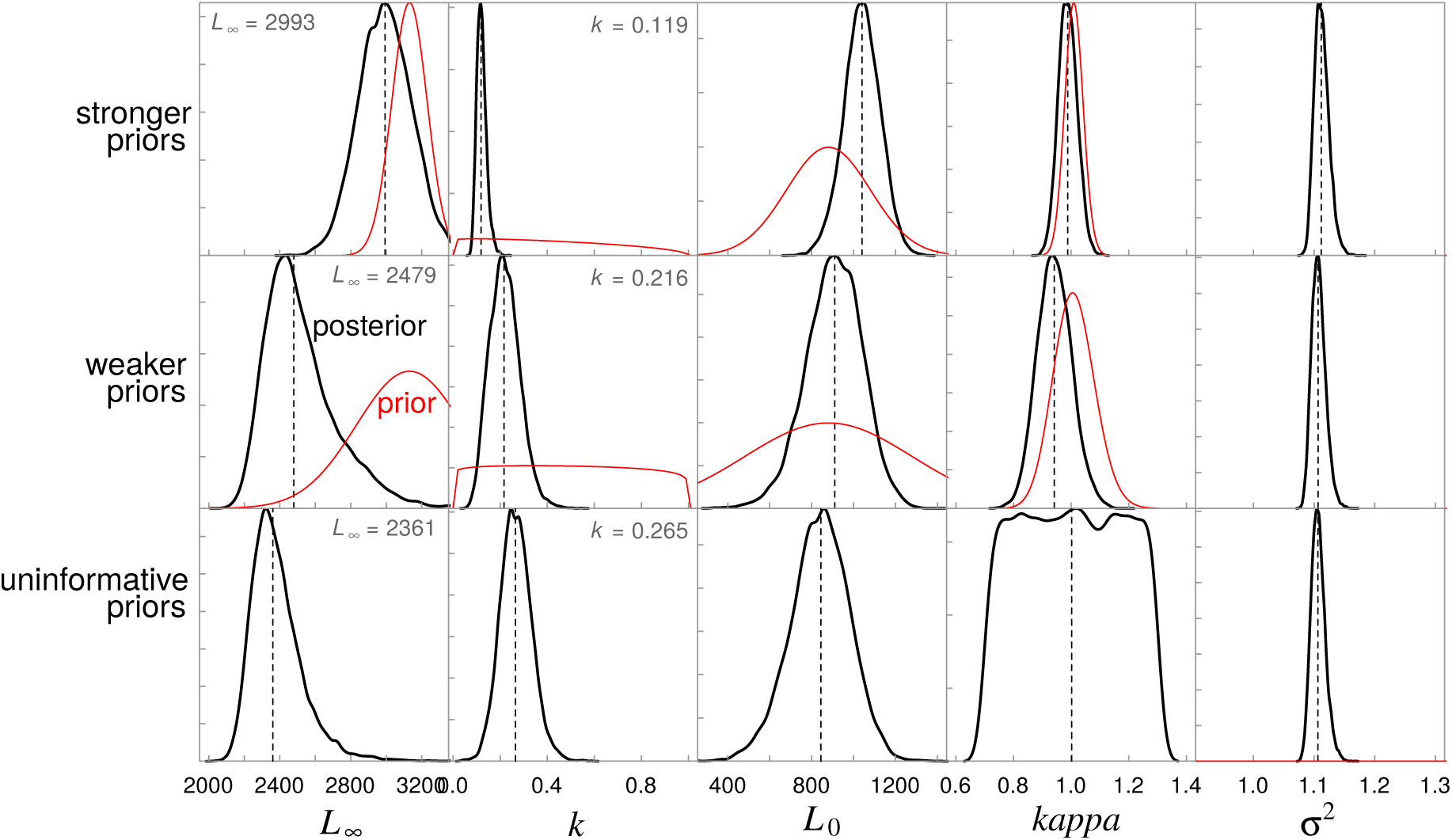
Prior and posterior distributions for the Spinetail devil Ray (*Mobula japanica*) von Bertalanffy growth parameters (*L*_∞_, *k*, and *L*_0_) and the error term (*σ*^2^) for the three Bayesian models with strong, weaker, and uninformative priors. Median values are shown by the dashed lines, posterior distributions by the black lines, and prior distributions by the red lines. No prior distributions are shown when priors are uninformative (uniform distribution).

**Figure 2:**
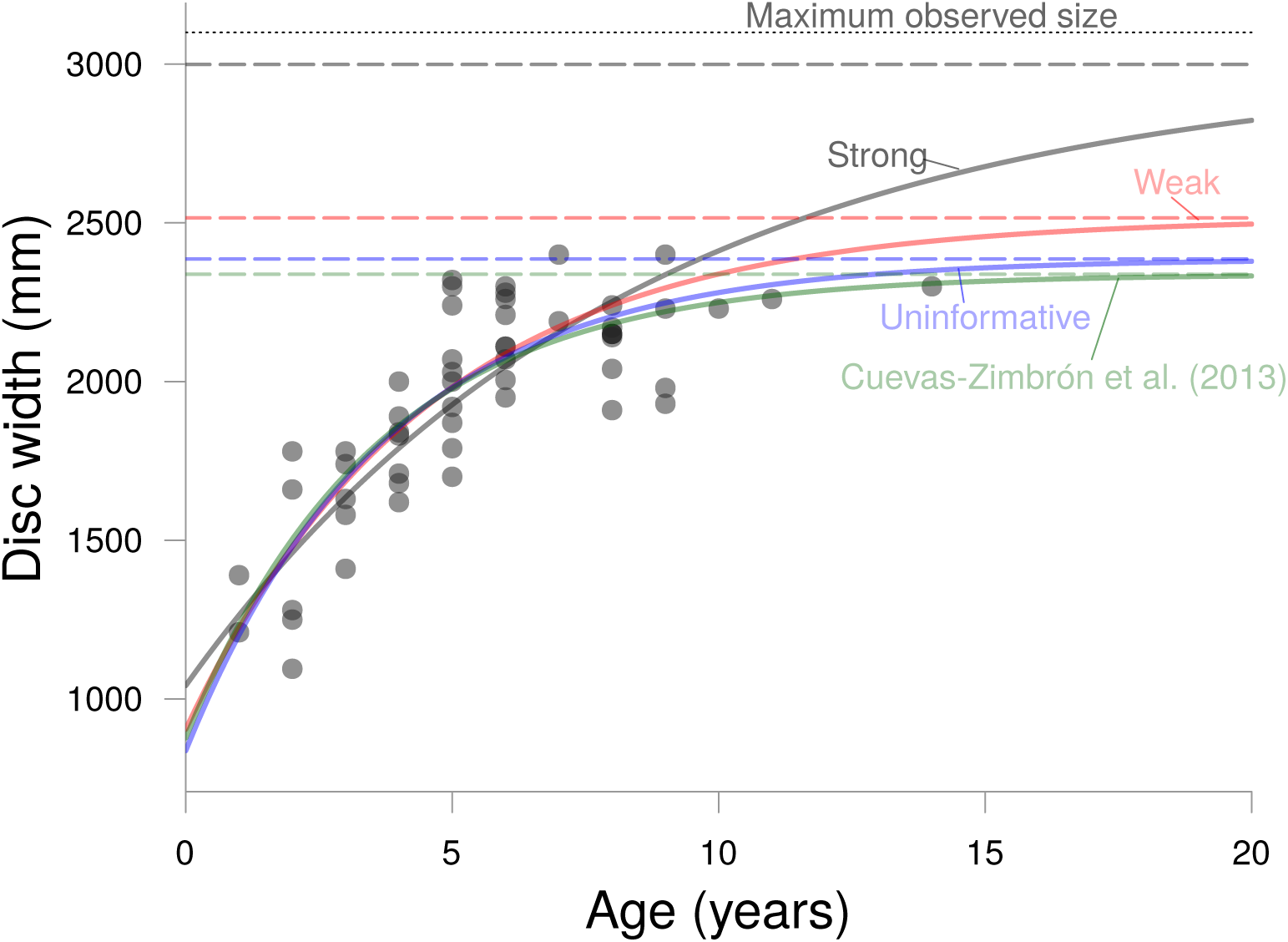
Length-at-age data for the Spinetail Devil Ray *Mobula japanica* showing the Bayesian von Bertalanffy growth curve fits for models with strong (grey), weaker (red), and uninformative (blue) priors, as well as the original model fit from Cuevas-Zimbrón et al. (2013). Dashed lines show the asymptotic size (*L*_∞_) estimates for each model. Dotted line represents the maximum known size for the species.

### Part 2. Estimating total mortality using the catch curve

Our catch curve analysis yielded a median estimate of *Z* = 0.254 year^−1^, with 95% of bootstrapped estimates ranging between 0.210 and 0.384 year^−1^ (Fig. 3). There were no devil rays aged 12 or 13, and therefore these age-classes were removed from the catch curve analysis before fitting each regression. Because these age classes are some of the oldest, removing these points is likely to provide more conservative estimates of total mortality. As a reference, we also ran the models without removing these age-classes but adding one to the number of individuals in age-class and got a very similar *Z* estimate (≈ 0.23 year^−1^). Assuming *Z* is approximately 0.25 year^−1^ is therefore a relatively conservative estimate of total mortality; we infer natural mortality *M* of *M. japanica* must be less than 0.25 year^−1^ or that 22.1% of the population was killed each year from a combination of natural and fishing mortality.

**Figure 3:**
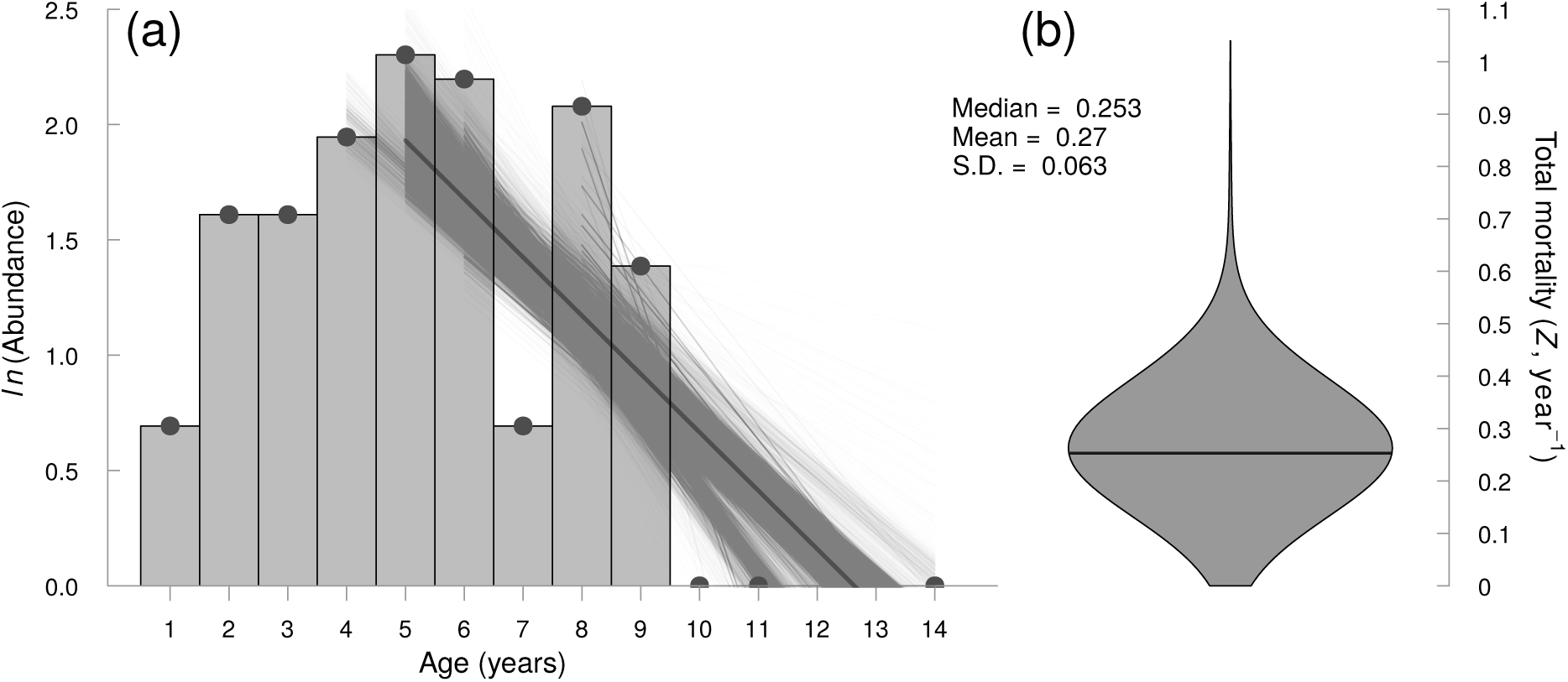
Estimation of total mortality *Z* from bootstrapped catch composition data for the Spinetail Devil Ray (*Mobula japanica*) from Cuevas-Zimbrón et al. (2013). (a) Catch curve of natural log abundance at age. The regression lines represent the estimated slopes when omitting different age-class subsets (as shown by the horizontal extent of each line), and resampling 80% of the data. Note that individual estimates of *Z* differ in the number of age-classes included for its computation, resulting in regression lines of different lengths. (b) Violin plot of estimated total mortality (*Z*) values calculated using the bootstrap resampling method. Estimates from different age-classes suggest an estimate of *Z* ≈ 0.25 year^−1^. The median is shown by the dark grey line.

### Part 3. Maximum population growth rate *r*_*max*_ of the Spinetail Devil Ray

From Part 1, we estimated that maximum lifespan was between 15 and 20 years. Combining this with estimated age at maturity, the median estimates of average lifespan for the Spinetail Devil Ray was 11.5 years, and therefore the median natural mortality *M* estimate was 0.087 year^−1^. From Part 2, we found that mortality *M* had an upper bound of 0.25 year^−1^(assuming *Z* = *M*). Using this information to create a bounded distribution for natural mortality in equation 10, we found the median maximum intrinsic rate of population increase *r*_*max*_ for devil rays is 0.077 year^−1^ (95th percentile = 0.042–0.108).

### Part 4. Comparing *Mobula r*_*max*_ to other chondrichthyans

Devil and manta rays have low intrinsic rate of population increase relative to other chondrichthyan species (Fig. 4). Among species with similar somatic growth rates, the Spinetail Devil Ray has the lowest *r*_*max*_ value (black diamond in Fig. 4a). This contrast is strongest when excluding deep-water chondrichthyans (white circles in Fig. 4), which tend to have much lower rates of population increase than shallow-water ones [33]. Our estimation of *r*_*max*_ for manta rays (grey diamond in Fig. 4) are comparable with our estimates for the Spinetail Devil Ray, albeit slightly lower (median of 0.068 year^−1^, 95th percentile = 0.045–0.088). Values of *r*_*max*_ for other large planktivorous elasmobranchs (Whale and Basking Sharks) are relatively high compared to manta and devil rays.

**Figure 4:**
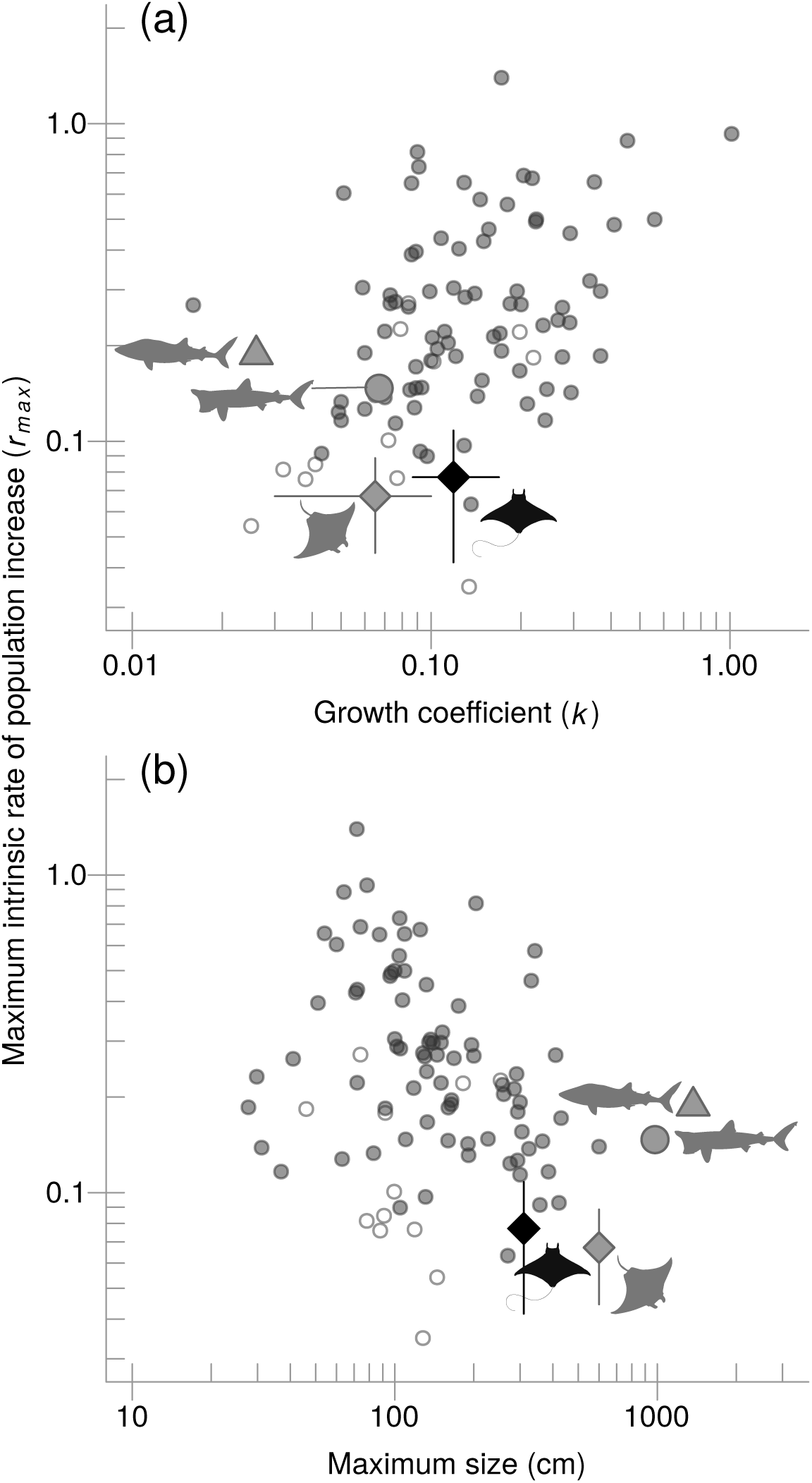
Comparison of maximum intrinsic rate of population increase (*r*_*max*_) for 96 elasmo-branch species arranged by (a) growth coefficient *k* and (b) maximum size. Small open circles represent deep sea species while small grey circles denote oceanic and shelf species. Four species are highlighted using silhouettes and larger symbols: the Spinetail Devil Ray (*Mobula japanica*) is shown by the black diamond, while the manta ray (*Manta spp*.), Whale Shark (*Rhincodon typus*) and Basking Shark (*Cetorhinus maximus*) are represented by the grey diamond, triangle and circle, respectively.

## Discussion

In this study, we examined multiple lines of evidence that suggest devil rays have relatively low productivity, and hence high risk of extinction compared to other chondrichthyans. The *r*_*max*_ of the Spinetail Devil Ray is comparable to that of manta rays, and much lower than that of other large planktivorous shallow-water chondrichthyans such as the Whale Shark and the Basking Shark (Fig. 4). We conclude the comparable extinction risks of devil and manta rays, coupled with the ongoing demand for their gill plates, suggest that conferring a similar degree of protection to all mobulids is warranted.

Next we consider three key questions that arise from our analyses: (1) Why do mobulid rays have such low productivity? (2) Can growth estimates be improved with prior knowledge? (3) Can small-scale fisheries cause local declines in devil ray populations?

### Why do mobulid rays have such low productivity?

We found that the Spinetail Devil Ray has similar productivity to manta rays, despite this species being half the size of the manta rays. This result suggests that smaller devil ray species are also likely to have very low productivity, probably due to their very low reproductive rates. Mobulid rays have at most a single pup annually or even biennially, while the Whale Shark can have litter sizes of up to 300 pups [34], therefore increasing the potential ability of this species to replenish its populations (notwithstanding differences in juvenile mortality). The Basking Shark has a litter size of six pups, which partly explains why its *r*_*max*_ is intermediate between mobulids and the Whale Shark. While mobulids mature relatively later with respect to their total lifespan than Basking Sharks and relatively earlier than Whale Sharks, they have lower lifetime fecundity than both Whale and Basking Sharks, limiting their productivity.

Our results are consistent with the correlation between low somatic growth rates, later maturation, large sizes, and elevated extinction risk that has been found in other marine fishes [35,36] (Fig. 4a). For example, in tunas and their relatives, somatic growth rate is the best predictor of overfishing, such that species with slower growth are more likely to be overfished as fishing mortality increases than species with faster growth [37], likely because the species that grow faster mature earlier.

### Can growth estimates be improved with prior knowledge?

Our method for estimating growth rate for the Spinetail Devil Ray provided lower estimates of the growth coefficient *k* than was reported in the original study [13], especially when we used strongly informative priors. When using strong priors the estimated asymptotic size was very close to the expectation of it being 90% of maximum size. On the other hand, our scenario with uninformative priors provided growth coefficient *k* estimates that are very similar to the original estimates, which were obtained by nonlinear least squares minimization (Fig. 2b). Our growth estimates from the model with strong priors are consistent with our expected values of *k* if length-at-age data were available for larger individuals. Given that the length-at-age data available only includes individuals up to two-thirds of the maximum size recorded for *M. japanica*, we believe that our approach provides more plausible estimates of growth rates when data are sparse. Our approach provides further evidence that Bayesian estimation is useful for data-sparse species as the available life history information can be easily incorporated in the form of prior distributions, particularly when missing samples of the largest or smallest individuals [15]. Incorporating prior information when fitting growth curves is an alternative to fixing model parameters, which often biases growth estimates [38]. In other words, using Bayesian inference allows us to incorporate out-of-sample knowledge of observed maximum sizes and sizes at birth, thus improving our estimates of growth rates and asymptotic size [15].

### Can small-scale fisheries cause local declines in devil ray populations?

The estimate of fishing mortality we calculated from the catch curve (*Z* – *M* = *F* = 0.163 year^−1^) is twice as high than our estimate of *r*_*max*_, which also represents the fishing mortality *F* expected to drive this species to extinction (*F*_*ext*_ = 0.077 year^−1^) [28]. Even though our estimate of fishing mortality is highly uncertain (Fig. 3b), the large discrepancy between our estimates of *F* and *F*_*ext*_ suggests that even if we are overestimating fishing mortality it was likely greater that *F*_*ext*_. Hence we infer that before the fishery ceased in 2007, the Spinetail Devil Ray population we examined was probably being fished unsustainably at a rate high enough to lead to eventual local extinction. Many teleost fisheries support fishing mortalities that are many times larger than natural mortality because of strong density dependence. However, mobulids likely have low capacity to compensate for fishing, because their large offspring and low fecundity suggest weak density-dependent regulation of populations [39,40].

The major caveats of using a catch curve analysis to estimate total mortality are that it assumes there is no size selectivity in catch, recruitment is constant, the population is closed, and that the catch is a large enough sample to sufficiently represent population age structure. These assumptions are also required in age and growth studies when using length-at-age data. Thus, our approach of estimating fishing mortality could be applied to other chondrichthyan growth studies, assuming that fishing is not systematically size selective and that metapopulation dynamics are not influencing the sample. Whether or not this latter assumption is valid for highly migratory elasmobranch species has yet to be tested.

Unregulated small-scale artisanal fisheries are targeting mobulids throughout the world [4, 5]. Our findings imply that there is little room for unmanaged artisanal fisheries to support sustainable international exports of gill plates or even domestic meat markets. Furthermore, the unsustainable fishing mortality stemming from the removal of relatively few individuals by an artisanal fishery suggests we urgently need to understand the consequences of bycatch of mobulid rays in industrial trawl, long line and purse seine fisheries [4,8]. The combination of high catch rates and low post-release survival suggest fishing mortality rates need to be understood and potentially minimized to ensure the future persistence of these species.

## Acknowledgements

We thank the fisherman of the fishing camp “Punta Arenas de la Ventana”, Baja California Sur, Mexico, for allowing collection of specimens and biological material. Field and lab work was supported by F. Galvan, N. Serrano, I. Mendez, A. Medellin, L. Castillo and C. Rodriguez, E. Diiaz, J. M. Alfaro and E. Bravo. We thank S. C. Anderson for statistical advice.

The original fieldwork was supported by the project “Historia Natural, Movimientos, Pesqueria y Criaderos, Administration de Mantas Mobulidas en el Golfo de California” of the Monterey Bay Aquarium. This research was funded by the J. Abbott/M. Fretwell Graduate Fellowship in Fisheries Biology (SAP), NSERC Discovery and Accelerator Grants (NKD), a Canada Research Chair (NKD), an NSF Postdoctoral Fellowship in Math and Biology (HKK; DBI-1305929), and grants to NKD from the John D. and Catherine T. MacArthur Foundation, the Leonardo DiCaprio Foundation, Disney Conservation Fund, and the Wildlife Conservation Society.

**Supporting Information**

